# Reproducible stool metagenomic biomarkers linked to the melanoma immunotherapy positive outcome

**DOI:** 10.1101/2022.04.01.486538

**Authors:** Evgenii I. Olekhnovich, Artem B. Ivanov, Anna A. Babkina, Arseniy A. Sokolov, Vladimir I. Ulyantsev, Dmitry E. Fedorov, Elena N. Ilina

## Abstract

The human gut microbiome plays an important role both in human’s health and disease. Recent studies have shown the undeniable influence of gut microbiota composition on cancer immunotherapy efficacy. However, these researches show a lack of consensus in defining reproducible metagenomic markers for a positive immunotherapy outcome. Accordingly, extended published data re-analysis may help reveal clearer associations between the composition of the gut microbiota and treatment response. In this study, we analyzed 358 stool metagenomes from 5 studies published earlier: 210 metagenomes from melanoma patients with positive immunotherapy outcome, 148 metagenomes from melanoma patients with negative immunotherapy outcome. The biomarkers were selected by the group comparison of patients’ stool samples with different treatment responses (47 responders vs 55 non-responders, 102 metagenomes). Selected biomarkers were verified using the available data describing the influence of the fecal microbiota transplantation on melanoma immunotherapy outcomes (9 donors, 6 responders, 19 non-responders, 256 metagenomes). According to our analysis, the resulting cross-study reproducible taxonomic biomarkers correspond to 12 Firmicutes, 4 Bacteroidetes, and 3 Actinobacteria. 140 gene groups were identified as reproducible functional biomarkers, including those potentially involved in production of immune-stimulating molecules and metabolites. In addition, we ranked taxonomic biomarkers by the number of functional biomarkers found in their metagenomic context. In other words, we predicted a list of the potential “most beneficial” bacteria for a positive response to melanoma immunotherapy. The obtained results can be used to make recommendations for the gut microbiota correction in cancer immunotherapy, and the resulting list of biomarkers can be considered for potential diagnostic ways for predicting melanoma immunotherapy outcome. Another important point is the functional biomarkers of positive immunotherapy outcome are distributed in different bacterial species that can explain the lack of consensus of defining melanoma immunotherapy beneficial species between different studies.

## Background

Until recently, cancer therapy was mainly focused on treatments such as surgery, ionizing radiation, and chemotherapy. However, lately, outstanding successes have been achieved in the treatment of melanoma and other cancerous types using methods of activating the patient’s own immunity - immunotherapy. This effect is provided by immune checkpoint inhibitors (ICIs), the administration of which induces T-lymphocyte-mediated immune responses against tumors [Borghaei et al., 2015]. However, responses to these treatments are often heterogeneous and non-durable. A significant number of patients fail to benefit from ICIs [Robert et al., 2019], while others have severe autoimmune side effects [Horvat et al., 2015].

The gut microbiota is well known to be involved in metabolic reactions that are important for human health. An active study of the human intestinal microbiome has shown a relationship with several clinical conditions including response to anticancer therapy with ICIs. Some studies using human biosamples describe differences in the microbiota composition of cancer patients with various responses to immunotherapy [Chaput et al., 2017; Frankel et al., 2017; Routhy et al., 2018; Matson et al., 2018; Gopalakrishnan et al., 2018; Peters et al., 2019; Spencer et al., 2021]. Moreover, transferring feces from a responder patient may modulate the outcome of a non-responder [Davar et al., 2021; Baruch et al., 2021]. These experiments confirm the vital role of the gut microbiota in regulating the response to cancer immunotherapy. Expansion of research in this area may lead not only to the creation of new diagnostic tools, but also, possibly, new medications and treatment strategies, that increase the effectiveness of immunotherapy. However, despite a large number of studies, researchers still cannot reach a consensus on gut microbial determinants of immunotherapy positive outcome. Moreover, performed meta-analyses [Limeta et al., 2020; Lee et al., 2022] also don’t clearly answer this question. Certainly, objective constant factors such as the general metagenomic data complexity and technical and/or biological variations, as well as specific individual factors, can cause difficulties. However, the choice of data analysis strategy may help to reveal previously unidentified trends and dependencies.

In the present work, we identify reproducible biomarkers linked to positive ICIs outcome using the compositional data analysis methods [Gloor et al., 2017] and stool metagenomes data from earlier published studies. We also verified our findings using the available data describing the influence of the fecal microbiota transplantation on melanoma immunotherapy outcomes. Additionally, we determined the connections between taxonomic and functional biomarkers that predicted potential “most beneficial” bacteria for a positive response to immunotherapy. Produced connections may explain the lack of consensus of identified taxonomic biomarkers between different studies.

## Results

### Discovery of reproducible taxonomic biomarkers linked to positive immunotherapy outcome

In total, 358 stool metagenomes from 5 metagenomic studies from the NCBI database were used in the analysis: 210 metagenomes from melanoma patients with positive immunotherapy outcome (responders, R group), 148 metagenomes from melanoma patients with negative immunotherapy outcome (non-responders, NR group). The group comparison of patients stool samples with different immunotherapy outcomes (Frankel 2017, Gopalakrishnan 2018, and Matson 2018: Group 1 datasets) as well as data analysis results of the fecal microbiota transplantation influence to melanoma immunotherapy outcome (Baruch 2021 and Davar 2021: Group 2 datasets) were used in the analysis. Sequencing statistics and other metadata about the metagenomes are presented in Supplementary Table S1. Discovery of reproducible taxonomic biomarkers linked to the positive immunotherapy outcome was performed in two stages. At the first stage, taxonomic biomarkers that distinguished melanoma patients by immunotherapy outcome were obtained using the differential rankings via Songbird approach individually for the Group 1 datasets. Log-ratio assessment of the selected features across metagenomes shows the clear statistically significant difference between R and NR groups (see Figure S1). Concatenate Songbird-derived rankings and differentials for selected species presented in Supplementary Figure 1.

Based on obtained results, Nine bacterial species (*Ruminococcus bromii, Blautia wexlerae, Bacteroides ovatus, Ruminococcus bicirculans, Barnesiella intestinihominis, Roseburia hominis, Alistipes putredinis, Bacteroides vulgatus* and *Roseburia faecis*) were positive treatment response predictors in at least two Group 1 datasets, while *Faecalibacterium prausnitzii* and *Eubacterium siraeum* in all Group 1 datasets.

However, the Group 2 datasets were processed in a special way. Using the RECAST approach, the donor-derived microbial diversity across the recipients metagenomes was identified while the Songbird was used for detecting donor-derived microbes which was linked to a immunotherapy positive outcome. A total of 102 donor-derived bacterial species were identified at least one recipient. The list of top 5 bacteria that colonize the most recipients includes *Eubacterium rectale, F. prausnitzii, A. putredinis, R. faecis*, and *Bacteroides uniformis* (see Figure S2). According to the analysis of variance, the donor-derived microbes content are more significantly linked to the donor subject in comparison to the positive immunotherapy outcome in both FMT datasets (PERMANOVA, R^2^ = 0.20, adj. p < 0.05; R^2^ = 0.07, adj. p < 0.05 respectively for Baruch 2021; R^2^ = 0.14, adj. p < 0.001; R^2^ = 0.09, adj. p < 0.001 for Davar 2021; Aitchison distance, 10000 permutations). Log-ratio assessment of the Songbird differential rankings features across donor-derived microbial profiles shows the clear statistically significant difference between R and NR groups (see Figure S3).

Reproducible biomarkers list in Group 2 datasets included *E. reclate, Acidaminococcus intestini, Collinsella aerofaciens, Roseburia intestinalis, R. faecis* and *F. prausnitzii*. However, only *Prevotella copri* is a reproducible negative immunotherapy predictor (see Figure 2). Interestingly, *E. rectale* engraftment is a more powerful predictor of positive treatment results for FMT-related datasets in comparison to the *F. prausnitzii*. Firmicutes were beneficial in both datasets (maintaining difference in the species content) while Bacteroidetes and Actinobacteria demonstrate contradictory behavior in the impact on immunotherapy context. For example, only in the Davar 2021 dataset *Bacteroides dorei* and *B. uniformis* were strongly linked to immunotherapy benefit but not in the Baruch 2021. Similarly, *A. putredinis* were linked to immunotherapy benefit in the Baruch 2021 dataset but not in the Davar 2021. Moreover, Actinobacteria colonization was only identified in the Baruch 2021 dataset as a significant predictor, while the main part Bacteroidetes only in the Davar 2021.

**Figure 1.**
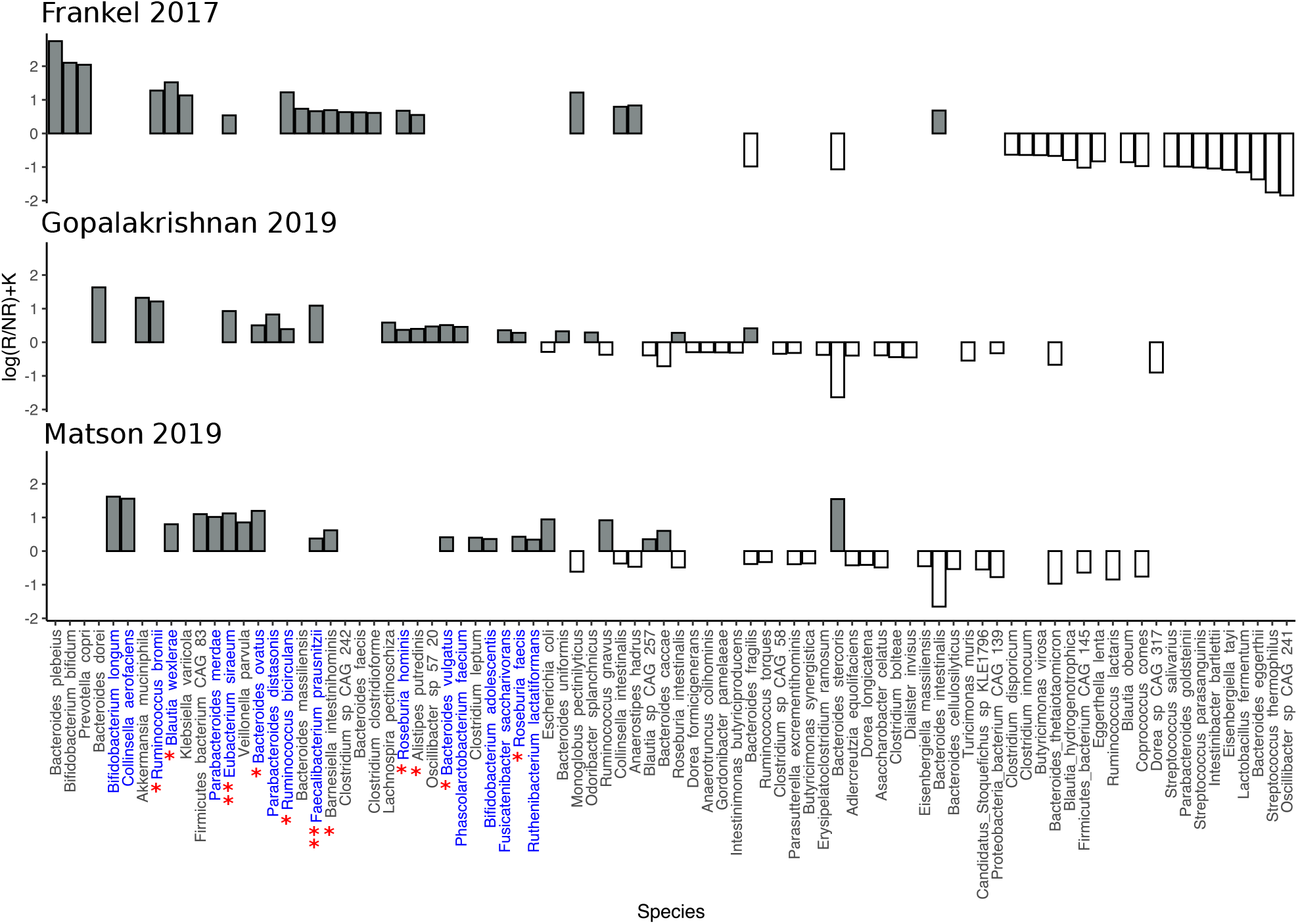
Species rankings and differentials derived from Songbird. X-axis shows microbial species, Y-axis shows differentials, which describe the log-fold change in features with respect to immunotherapy outcome. Positive coefficient levels correspond to positive associations. The red star denoted species with positive associations in two datasets while two red stars - in three datasets. Species that have been added to the list of reproducible biomarkers are highlighted in blue.

**Figure 2.**
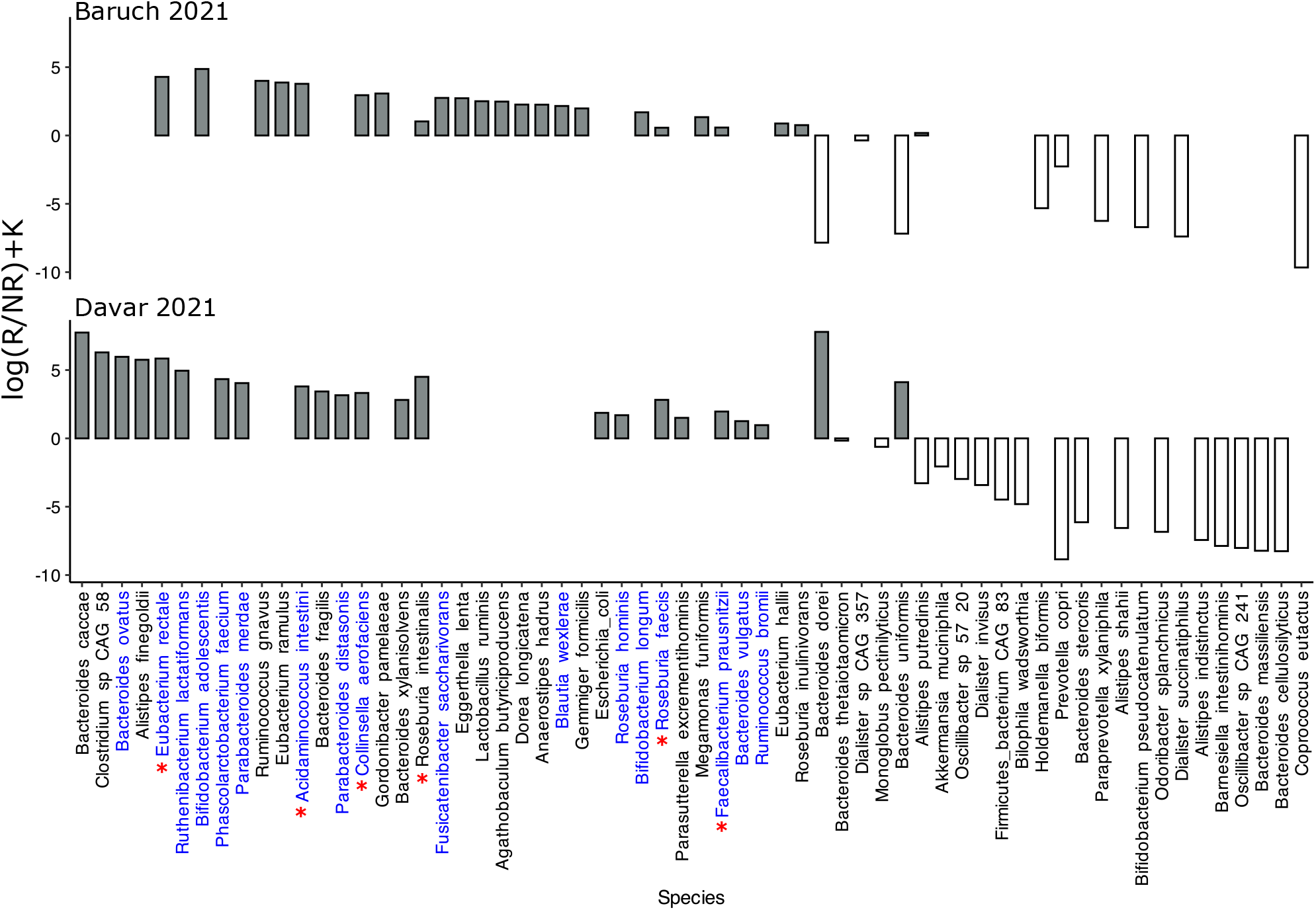
Donor-derived species rankings and differentials derived from Songbird. X-axis shows microbial species, Y-axis shows differentials, which describe the log-fold change in features with respect to immunotherapy outcome. Positive coefficient levels correspond to immunotherapy response. The red stars denoted species with positive associations in both Baruch 2021 and Davar 2021 datasets. Species that have been added to the list of reproducible biomarkers are highlighted in blue.

It is clear that the FMT useful effect is strongly dependent on both donor subject and donor specific microbial content. However, not all recipients have benefited from the impact of a successful donor.

The second stage of analysis was the formation of a list of reproducible biomarkers using the following rules: 1) microbial species were added to the list which linked to positive immunotherapy outcome in more than one dataset; 2) microbial species that were classified as linked to negative immunotherapy outcome in at least one dataset were excluded from biomarkers list regardless of the number of datasets in which it were associated with a positive outcome. For additional validation of the identified taxonomic biomarkers, we cite publications that also demonstrate the links to cancer immunotherapy benefit and added that to the resulting table. The obtained results presented in Table 1.

**Table 1.** Reproducible taxonomic biomarkers associated with immunotherapy positive outcome.

| Species | Phyla | Analysis | Cites |
| --- | --- | --- | --- |
| *Faecalibacterium prausnitzii* | Firmicutes | Frankel et al., 2017; Gopalakrishnan et al., 2018; Matson et al., 2018; Baruch et al., 2021; Davar et al., 2021; | Chaput et al., 2017; Frankel et al., 2017; Gopalakrishnan et al., 2018; Matson et al., 2018; Peters et al., 2019; Coutzac et al., 2020; Limeta et al., 2020; Mao et al., 2021; Spencer et al., 2021; |
| *Roseburia faecis* | Firmicutes | Gopalakrishnan et al., 2018; Matson et al., 2018; Baruch et al., 2021; Davar et al., 2021; | Lee et al., 2022; |
| *Bacteroides ovatus* | Bacteroidetes | Gopalakrishnan et al., 2018; Matson et al., 2018; Davar et al., 2021; | Heshiki et al., 2020; |
| *Bacteroides vulgatus* | Bacteroidetes | Gopalakrishnan et al., 2018; Matson et al., 2018; Davar et al., 2021; | Usyk et al., 2021; Huang et al., 2021; |
| *Blautia wexlerae* | Firmicutes | Frankel et al., 2017; Matson et al., 2019; Baruch et al., 2021; |  |
| *Collinsella aerofaciens* | Actinobacteria | Matson et al., 2018; Baruch et al., 2021; Davar et al., 2021; | Matson et al., 2018; |
| *Eubacterium siraeum* | Firmicutes | Frankel et al., 2017; Gopalakrishnan et al., 2018; Matson et al., 2018; | Derosa et al., 2020; Mao et al., 2021; |
| *Roseburia hominis* | Firmicutes | Frankel et al., 2017; Gopalakrishnan et al., 2018; Davar et al., 2021; | Lee et al., 2022; |
| *Ruminococcus bromii* | Firmicutes | Frankel et al., 2017; Gopalakrishnan et al., 2018; Davar et al., 2021; | Gopalakrishnan et al., 2018; Zheng et al., 2019; Fedorov et al., 2020; |
| *Acidaminococcus intestini* | Firmicutes | Baruch et al., 2021; Davar et al., 2021; |  |
| *Bifidobacterium adolescentis* | Actinobacteria | Matson et al., 2019; Baruch et al., 2021 | Sivan et al., 2015; Matson et al., 2018; Fedorov et al., 2020; Sun et al., 2020; |
| *Bifidobacterium longum* | Actinobacteria | Matson et al., 2019; Baruch et al., 2021 | Sivan et al., 2015; Matson et al., 2018; Jin et al., 2019; Sun et al., 2020; |
| *Eubacterium rectale* | Firmicutes | Baruch et al., 2021; Davar et al., 2021; |  |
| *Fusicatenibacter saccharivorans* | Firmicutes | Gopalakrishnan et al., 2018; Baruch et al., 2021; |  |
| *Parabacteroides distasonis* | Bacteroidetes | Gopalakrishnan et al., 2018; Davar et al., 2021; | Tanoue et al., 2019; Huang et al., 2021; |
| *Parabacteroides merdae* | Bacteroidetes | Matson et al., 2018; Davar et al., 2021; | Matson et al., 2018; Fedorov et al., 2020; |
| *Phascolarctobacterium faecium* | Firmicutes | Gopalakrishnan et al., 2018; Davar et al., 2021; |  |
| *Ruminococcus bicirculans* | Firmicutes | Frankel et al., 2017;<br>Gopalakrishnan et al.,<br>2018; |  |
| *Ruthenibacterium lactatiformans* | Firmicutes | Matson et al., 2018; Davar<br>et al., 2021; | Tanoue et al., 2019; |

The list of reproducible taxonomic biomarkers linked to positive immunotherapy outcomes included 19 bacterial species: 12 Firmicutes, 4 Bacteroides and 3 Actinobacteria. Many major short-chain fatty acid producers such as *F. prausnitzii, R. faecis, R. hominis, R. bromii*, and *Eubacterium rectale* have been included to the reproducible biomarkers list. However, *F. prausnitzii* was identified as a significant predictor of positive immunotherapy results in all datasets added to the analysis. In addition, we found 9 studies in the scientific literature that also note the beneficial role of *F. prausnitzii* in immunotherapy response. Interestingly, all defined reproducible taxonomic biomarkers were identified as colonizers in FMT-datasets (see Figure S2). However, only 17 out of 19 (excluding *Ruminococcus bicirculans* and *E. siraeum*) were defined as positive immunotherapy in FMT-related datasets. Moreover, *E. reclate* and *A. intestini* were defined as positive immunotherapy outcome predictors only in FMT-related datasets. It should be additionally noted that *F. prausnitzii* and *Roseburia* spp. have been identified as cross-cohort biomarkers of response to ICIs therapy in previously published meta-analyses [Limeta et al., 2020; Lee et al., 2022] which further confirms the role of these bacteria in the positive ICIs response.

### Discovery of reproducible functional biomarkers linked to positive immunotherapy outcome

Discovery of reproducible functional biomarkers linked to a positive immunotherapy outcome was performed in an identical way to taxonomic characteristics. Log-ratio assessment of the selected KOG across studied metagenomes or donor-derived microbial profiles for added datasets shows the clear statistical significant difference between R and NR groups (Wilcoxon rank-sum test, p < 0.001). A total of 140 KOGs linked to positive immunotherapy outcome in more than one dataset were identified (see Supplementary Table S2).

Among the list of reproducible functional biomarkers are KOG involved in the polysaccharides metabolism (K16148, K16147, K01210, K01218, K01136), peptidoglycan (K07284, K05364, K12554) and lipopolysaccharide (K12983, K12997) biosynthesis as well as fatty acids biosynthesis (K11533, K11263). It should be noted that gluABCD glutamate uptake system (K10005, K10006, K10007, K10008) and gfrABCD PTS system (K19506, K19507, K19508, K19509) of Maillard reaction products (MRPs) fructoselysine/glucoselysine utilization are also predictors of positive immunotherapy outcome. Among other things, significant predictors of response were gene groups involved in the biosynthesis of cofactors such as cobalamin (K13542, K24866) and ubiquinone (K03688). Moreover, the KOG corresponds to sporulation (K18955, K06383, K06297, K06313, K06330, K06413) and motility/secretion system (K02653, K02398, K02417, K02420, K02278) also were linked to the success of ICIs therapy outcome.

Using the MetaCharchant and Kraken2 tools, we studied the metagenomic context of selected reproducible functional biomarkers described above (see Supplementary Table S3). According to the analysis of variance using PERMANOVA, different bacterial phyla were linked to different functional content (R^2^ = 0.17, adj. p < 0.001; Bray-Curtis dissimilarity, 10000 permutations). More KOG-biomarkers were identified in Actinobacteria and Firmicutes, while Bacteroidetes were least linked to selected gene groups (Wilcoxon rank-sum test, p < 0.05). Top bacteria in which most of the functions linked to a positive response to cancer immunotherapy include *B. longum, B. adolescentis, E. rectale, F. prausnitzii* (see Figure 3A). Moreover, 12 out of 19 (∼63%) previously described reproducible taxonomic biomarkers were linked to KOG-biomarkers.

**Figure 3.**
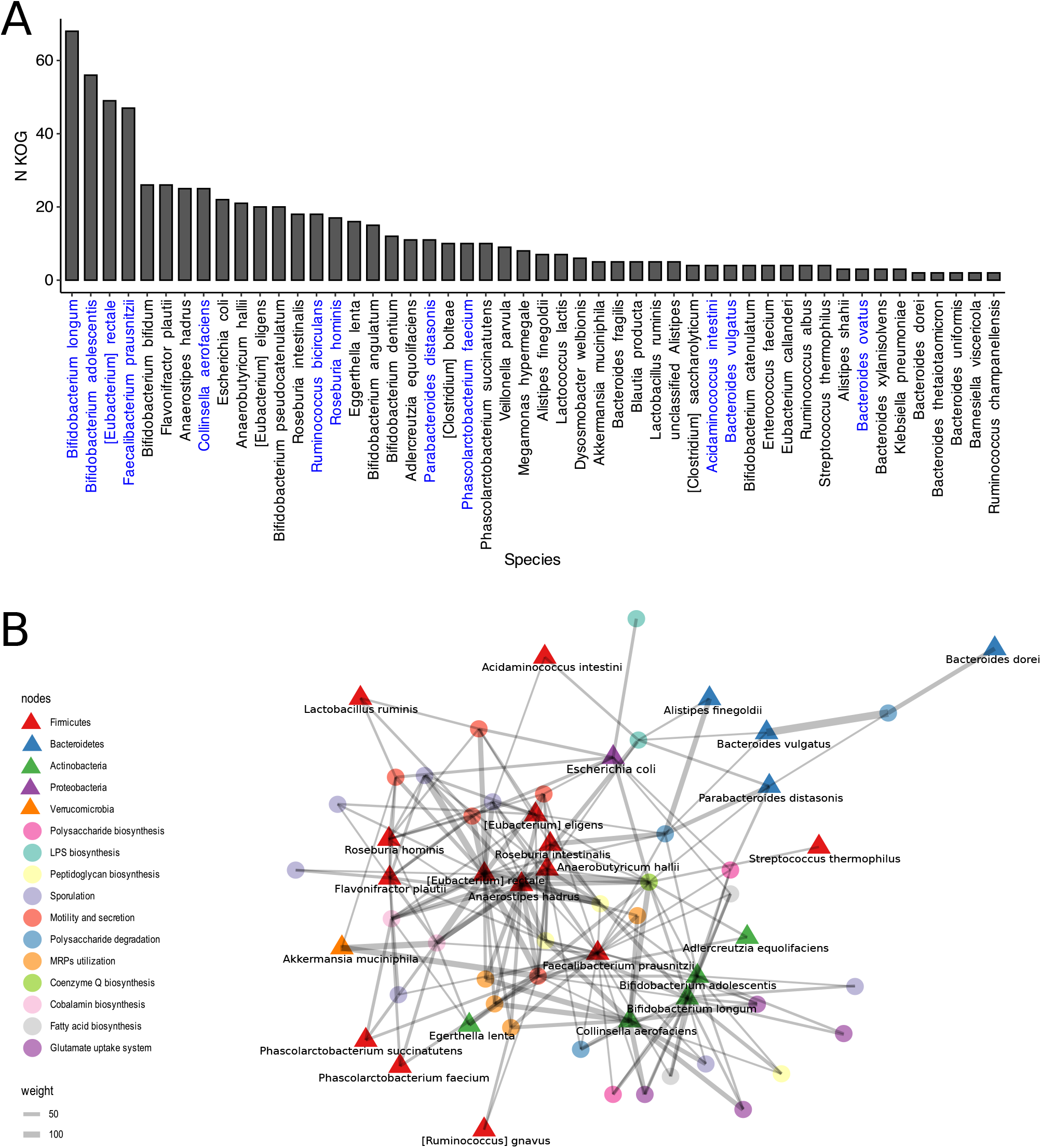
Links between bacterial species and functional reproducible biomarkers of immunotherapy positive outcome. **(A)**. The number of KOGs found in the metagenomic context of bacterial species. X-axis shows bacterial species as well as Y-axis - number of KOGs. Species that have been added to the list of reproducible biomarkers are highlighted in blue. **(B)** The network shows KOG groups links to bacterial species. Only KOGs that were presented in Table 2 were added to the analysis to avoid visual overload of figure. Triangles correspond to Bacterial phyla while circles to KOGs.

In addition, network analysis of the associations of selected functional biomarkers and bacterial species was performed (see Figure 3B). According to this analysis, bacterial phyla differed in the content of functional biomarkers also as demonstrated above. For example, cobalamin biosynthesis is strongly linked to the Firmicutes and Verrucomicrobia (*Akkermansia muciniphila*) while the glutamate uptake system to the Actinobacteria. However, ubiquinone biosynthesis and MRPs utilization system have been presented in both Actinobacteria and Firmicutes. It is worth noting that according to this analysis Bacteroidetes have been associated only with gene groups involved in polysaccharides degradation. Interestingly, *F. prausnitzii* occupies an intermediate position between Firmicutes and Actinobacteria clusters and is linked to the specific KOGs of both bacterial phyla. It is worth also noting that the list of species associated with functions predicted ICIs outcome included *F. prausnitzii, Akkermansia muciniphila*, and *Roseburia* spp. which was earlier determined as reproducible biomarkers in meta-analyses [Limeta et al., 2020; Lee et al., 2022].

### Discussion

The world scientific community is conducting extensive research to determine the influence of the human intestinal microbiota to positive outcomes of cancer immunotherapy. However, despite a large number of studies, researchers still cannot reach a consensus on the detection of reproducible gut microbial determinants of immunotherapy positive outcome. In this way, the development and application of new types of metagenomic analysis tools as well as published data re-analysis may help reveal new associations between the composition of the gut microbiota and immunotherapy outcome.

We analyzed the gut metagenomes of melanoma patients from five published studies and determined nineteen cross-study taxonomic biomarkers linked to positive immunotherapy outcome. This list included twelve Firmicutes, four Bacteroidetes, and three Actinobacteria species. However, only *F. prausnitzii* is reproducible for all datasets included in the analysis. Identified biomarkers are confirmed by many published studies [Sivan et al., 2015; Chaput et al., 2017; Frankel et al., 2017; Gopalakrishnan et al., 2018; Matson et al., 2018; Jin et al., 2019; Peters et al., 2019; Zheng et al., 2019; Tanoe et al., 2019; Derosa et al., 2020; Coutzac et al., 2020; Limeta et al., 2020; Fedorov et al., 2020; Heshiki et al., 2020; Sun et al., 2020; Lee et al., 2021; Spencer et al., 2021; Mao et al., 2021; Usyk et al., 2021; Lee et al., 2022]. Additionally, we identified 140 KOGs as reproducible functional biomarkers of positive immunotherapy outcome corresponds to fatty acids biosynthesis, polysaccharide biosynthesis and degradation, peptidoglycan and lipopolysaccharide biosynthesis, cobalamin biosynthesis, fructoselysine/glucoselysine utilization system, sporulation, and motility/secretion system.

It is known that gut microbial community promotes large scale digestive metabolic function involved in health, some of which may be associated with improved immune system function and immunotherapy outcome. Identified biomarkers can be linked to published results on molecular mechanisms involved in response to immunotherapy. For example, increased intake of plant polysaccharides improves the outcome of immunotherapy [Huang et al., 2021; Spencer et al., 2021]. Moreover, a large number of taxonomic biomarkers are major butyrate and other short-chain fatty acid producers. It’s known that gut microbiome derived SCFAs are necessary for the implementation of normal body metabolism and modulate cytotoxic immunity, which improves immunotherapy efficiency [Luu et al., 2021; He et al., 2021; Dann et al., 2021]. On the other hand, gene groups involved in sporulation, motility and polysaccharides production were also identified as positive immunotherapy outcome biomarkers. These gene groups can be characterized as pattern of recognition receptors ligands, which involved to modulation of host immunity and immunotherapy effectiveness [Vétizou et al., 2015; Daillère et al., 2016; Pushalkar et al., 2018; Griffin et al., 2021; Lee et al., 2021].

Nevertheless, novel findings that may be important in modulating immunotherapy outcome by gut microbes can be described. Gut bacteria may affect food safety through Maillard reaction products utilization [Wolf et al., 2019]. We have determined that the gfrABCD PTS system is used by bacteria as a fructoselysine/glucosolysine disposal system to predict a positive outcome of immunotherapy. Moreover, fructoselysine for intestinal bacteria may be used as additional butyrate production substrate [Bui et al., 2015], which increases the value of this finding for interpreting the beneficial properties of the microbiota for cancer immunotherapy. In addition, we ranked bacterial species by the number of KOG biomarkers found in their metagenomic environment. The largest number of KOG biomarkers were found in the metagenomic context of four bacterial species such as *B. longum, B. adolescentis, E. rectale*, and *F. prausnitzii*. Based on these results, we hypothesize that the distribution of beneficial functions in different bacterial species is the main reason for the lack of consensus in taxonomic biomarkers between different studies. Moreover, *E. rectale, F. prausnitzii* and bifidobacteria were defined as bacteria with immunomodulatory potential in comparison of COVID-19 patients cohort and healthy persons [Yeoh et al., 2021].

Fecal microbiota transplantation impact to immunotherapy outcomes was strongly linked to donor variable. However, beneficial donors were not successful in improving treatment outcomes in all cases. In other words, the FMT immunomodulatory success may depend on both the donor stool characteristics and the recipient gut environment condition (or other unknown factors). Moreover, species that have been linked to therapy success have also colonized patients with negative outcomes. It’s possible that broad spectrum donor-derived species engraftment is a “side effect” of FMT [Goloshchapov et al., 2019], while special “hidden” donor microbiome characteristics such as bacteriophages, or specific strains-produced molecular structures can be linked to beneficial immunotherapy response [Ott et al., 2019; Lee et al., 2021]. On the other hand, microbial load (which cannot be assessed using metagenomic data) in donor stool and/or recipient mucosa [Sarrabayrouse et al., 2020] may play a decisive role for the FMT success.

## Conclusions

It is known that the gut microbiota has many positive effects on the host, ranging from metabolic function to the influence of microbial-derived molecular structures on specific cell receptors. Since metagenomic sequencing does not detect fecal microbial load, however, it can be assumed, a “general microbiome signal” detected by human cells is degraded in non-responders in comparison to responders. On the other hand, the special characteristics of the responders intestinal community which can be transferred from one person to another may determine the impact on the outcome of immunotherapy. Anyway, the true biological nature of the observed phenomena has yet to be established. However, some useful conclusions can already be drawn. For example, digestive fibers and probiotic bifidobacterium can be included in treatment regimens to improve immunotherapy outcome, while the question of choosing the most useful strains among commercial probiotics remains open yet.

## Methods

### Metagenomic data

Gut metagenomes of melanoma patients with different responses to immune checkpoint inhibitors were collected before any interventions from three published studies [Frankel et al., 2017; Gopalakrishnan et al., 2018; Matson et al., 2018]. These data were used for association analysis between taxonomic or functional metagenomic profiles and immunotherapy outcome. Additional data reflecting the impact of FMT to immunotherapy outcome [Baruch et al., 2021; Davar et al., 2021] was used for identified biomarkers verification. The features crossover between at least two datasets were considered reproducible biomarkers. In total, 358 metagenomic stool samples were used in the study, among them 210 metagenomes from melanoma patients with positive immunotherapy outcome (responders, R) group - R), 148 metagenomes from melanoma patients with negative outcome (non-responders, NR) group - NR).

### Data analysis strategy

Raw metagenomic data were downloaded from public NCBI/EBI repositories using fastq-dump from the SRA Toolkit [Sherry and Chunlin, 2012] while quality assessment was performed with FastQC [https://github.com/s-andrews/FastQC]. Technical sequences and low-quality bases were trimmed with the Trimmomatic [Bolger et al., 2014] with threshold for sequencing quality > 30. The human sequences were removed from metagenomes by bbmap [Bushnell, 2014] and GRCh37 human genome version. Described metagenomics reads preprocessing computational steps were implemented in the Assnake metagenomics pipeline [https://github.com/ASSNAKE]. Donor-derived reads of post-FMT stool metagenomes in FMT-related datasets were identified using the RECAST algorithm [Olekhnovich et al., 2021]. Taxonomic and functional profiles of processed stool metagenomes or donor-derived reads were obtained using MetaPhlAn3 [Beghini et al., 2021] and HUMAnN2 approaches [Franzosa et al., 2018] with and the KEGG database [Kanehisa et al., 2017].

Detection of reproducible taxonomic and functional biomarkers associated with a positive immunotherapy outcome was performed in two stages. At the first stage, using the taxonomic and functional profiles of stool metagenomes or donor-derived reads (for FMT-related data), increased and decreased microbial species were identified in patients with a positive outcome of immunotherapy using the Songbird [Morton et al., 2019] and Qurro [Fedarko et al., 2020] implemented in the QIIME2 framework [Bolyen et al., 2019]. The second stage was the formation of a list of reproducible biomarkers using the following rules: 1) microbial species were added to the list that were associated with a positive immunotherapy outcome in more than one dataset; 2) microbial species that were classified as negatively associated with positive immunotherapy outcome in at least one data set, regardless of the number of data sets in which it were associated with a positive outcome, were excluded from this list. The search of links between taxonomic and functional biomarkers was performed as follows. Reconstruction of the functional biomarkers metagenomic context was performed using MetaCherchant [Olekhnovich et al., 2018] while its taxonomic annotation was obtained by Kraken2 [Wood et al., 2019].

Additional statistical analysis and visualizations were carried out using *vegan* package [Oksanen et al., 2013], ggplot2 library and standard statistical procedures implemented in GNU/R. PERMANOVA (*adonis* function from the *vegan* package), Bray-Curtis dissimilarity [Oksanen et al., 2013] and Aitchison distance [Aitchison et al., 1992; Aitchison et al., 1997] were used as measures for comparing taxonomy profiles of reads categories and functional biomarkers content.

## Acknowledgments

We thank the Center for Precision Genome Editing and Genetic Technologies for Biomedicine, Federal Research and Clinical Center of Physical-Chemical Medicine of the Federal Medical Biological Agency for providing computational resources for this project.

## Authors’ contributions

**EIO** - performed computational experiments, performed metagenomic data analysis and visualization, interpretation of obtained results, contributed to research idea, and wrote the manuscript. **AAB, AAS** - metagenomic data analysis and visualization support. **ABI, VIU** - contributed to metagenomic data analysis and visualization, contributed to manuscript preparation. **DEF** - metagenomic data management, support of the bioinformatics computational protocols, contributed to research idea, contributed to interpretation of obtained results and manuscript preparation. **ENI** - research idea, contributed to interpretation of the obtained results and manuscript preparation.

## Availability of data and materials

The study used data from open sources, which are available at NCBI Sequence Read Archives under the BioProjects accession numbers PRJNA397906, PRJEB22893, PRJNA399742, PRJNA678737, PRJNA67286. All the results of the project were presented in the article text and in additional materials.

## Supplemental Materials

**Supplementary Table S1** - Summary data and sequencing statistics about metagenomes included in the analysis.

**Supplementary Table S2** - Reproducible functional biomarkers associated with immunotherapy positive outcome with differential coefficients.

**Supplementary Table S3** - Links between functional biomarkers and bacterial species.

**Supplementary Figure S1** - Feature rankings and differentials derived from Songbird (A, C, E) and boxplots of the log ratios of the selected features (B, D, F). The log-ratios statistical assessment shows a clear significant difference between R (responders) and NR (nonresponders) groups (Wilcoxon rank-sum test, p < 0.001 for Frankel 2017 and Matson 2018 and p < 0.05 for Gopalakrishnan 2018). Selected features (bacterial species) added for log ratios assessment denoted blue and green colors corresponding to R and NR groups, respectively.

**Supplementary Figure S2** - Donor-derived metagenomic reads per species found in the post-FMT recipients metagenomes. Different colors showed different immunotherapy outcome. The dot size corresponds to the number of metagenomic reads. Species that have been added to the list of reproducible biomarkers are highlighted in blue.

**Supplementary Figure S3** - Feature rankings and differentials derived from Songbird (A, C) and boxplots of the log ratios of the selected features (B, D). The log-ratios statistical assessment shows a clear significant difference in donor-derived microbial profiles between R (responders) and NR (non-responders) groups (Wilcoxon rank-sum test, p < 0.001). Selected features (bacterial species) added for log ratios assessment denoted blue and green colors corresponding to R and NR groups, respectively.

## References

1. Borghaei, Hossein, et al. “Nivolumab versus docetaxel in advanced nonsquamous non– small-cell lung cancer.” New England Journal of Medicine 373.17 (2015): 1627–1639.

2. Robert, Caroline, et al. “Pembrolizumab versus ipilimumab in advanced melanoma.” New England Journal of Medicine 372.26 (2015): 2521–2532.

3. Horvat, Troy Z., et al. “Immune-related adverse events, need for systemic immunosuppression, and effects on survival and time to treatment failure in patients with melanoma treated with ipilimumab at Memorial Sloan Kettering Cancer Center.” Journal of Clinical Oncology 33.28 (2015): 3193.

4. Chaput, Nathalie, et al. “Baseline gut microbiota predicts clinical response and colitis in metastatic melanoma patients treated with ipilimumab.” Annals of Oncology 28.6 (2017): 1368–1379.

5. Frankel, Arthur E., et al. “Metagenomic shotgun sequencing and unbiased metabolomic profiling identify specific human gut microbiota and metabolites associated with immune checkpoint therapy efficacy in melanoma patients.” Neoplasia 19.10 (2017): 848–855.

6. Gopalakrishnan, Vancheswaran, et al. “Gut microbiome modulates response to anti–PD-1 immunotherapy in melanoma patients.” Science 359.6371 (2018): 97–103.

7. Matson, Vyara, et al. “The commensal microbiome is associated with anti–PD-1 efficacy in metastatic melanoma patients.” Science 359.6371 (2018): 104–108.

8. Routy, Bertrand, et al. “Gut microbiome influences efficacy of PD-1–based immunotherapy against epithelial tumors.” Science 359.6371 (2018): 91–97.

9. Peters, Brandilyn A., et al. “Relating the gut metagenome and metatranscriptome to immunotherapy responses in melanoma patients.” Genome medicine 11.1 (2019): 1–14.

10. Spencer, Christine N., et al. “Dietary fiber and probiotics influence the gut microbiome and melanoma immunotherapy response.” Science 374.6575 (2021): 1632–1640.

11. Davar, Diwakar, et al. “Fecal microbiota transplant overcomes resistance to anti–PD-1 therapy in melanoma patients.” Science 371.6529 (2021): 595–602.

12. Baruch, Erez N., et al. “Fecal microbiota transplant promotes response in immunotherapy-refractory melanoma patients.” Science 371.6529 (2021): 602–609.

13. Limeta, Angelo, et al. “Meta-analysis of the gut microbiota in predicting response to cancer immunotherapy in metastatic melanoma.” JCI insight 5.23 (2020).

14. Lee, Karla A., et al. “Cross-cohort gut microbiome associations with immune checkpoint inhibitor response in advanced melanoma.” Nature Medicine (2022): 1–10.

15. Gloor, Gregory B., et al. “Microbiome datasets are compositional: and this is not optional.” Frontiers in microbiology 8 (2017): 2224.

16. Coutzac, Clélia, et al. “Systemic short chain fatty acids limit antitumor effect of CTLA-4 blockade in hosts with cancer.” Nature communications 11.1 (2020): 1–13.

17. Mao, Jinzhu, et al. “Gut microbiome is associated with the clinical response to anti-PD-1 based immunotherapy in hepatobiliary cancers.” Journal for Immunotherapy of Cancer 9.12 (2021).

18. Heshiki, Yoshitaro, et al. “Predictable modulation of cancer treatment outcomes by the gut microbiota.” Microbiome 8.1 (2020): 1–14.

19. Usyk, Mykhaylo, et al. “Bacteroides vulgatus and Bacteroides dorei predict immune-related adverse events in immune checkpoint blockade treatment of metastatic melanoma.” Genome medicine 13.1 (2021): 1–11.

20. Huang, Jumin, et al. “Ginseng polysaccharides alter the gut microbiota and kynurenine/tryptophan ratio, potentiating the antitumor effect of antiprogrammed cell death 1/programmed cell death ligand 1 (anti-PD-1/PD-L1) immunotherapy.” Gut (2021).

21. Luu, Maik, et al. “Microbial short-chain fatty acids modulate CD8+ T cell responses and improve adoptive immunotherapy for cancer.” Nature Communications 12.1 (2021): 1–12.

22. Danne, Camille, and Harry Sokol. “Butyrate, a new microbiota-dependent player in CD8+ T cells immunity and cancer therapy?.” Cell Reports Medicine 2.7 (2021): 100328.

23. He, Yao, et al. “Gut microbial metabolites facilitate anticancer therapy efficacy by modulating cytotoxic CD8+ T cell immunity.” Cell Metabolism 33.5 (2021): 988–1000.

24. Derosa, Lisa, et al. “Gut bacteria composition drives primary resistance to cancer immunotherapy in renal cell carcinoma patients.” European urology 78.2 (2020): 195–206.

25. Zheng, Yi, et al. “Gut microbiome affects the response to anti-PD-1 immunotherapy in patients with hepatocellular carcinoma.” Journal for immunotherapy of cancer 7.1 (2019): 1–7.

26. Fedorov, D. E., et al. “Gut Microbiome as a Predictor of the Anti-PD-1 Therapy Success: Metagenomic Data Analysis.” Biochemistry (Moscow), Supplement Series B: Biomedical Chemistry 15.2 (2021): 161–165.

27. Sivan, Ayelet, et al. “Commensal Bifidobacterium promotes antitumor immunity and facilitates anti–PD-L1 efficacy.” Science 350.6264 (2015): 1084–1089.

28. Sun, Shan, et al. “Bifidobacterium alters the gut microbiota and modulates the functional metabolism of T regulatory cells in the context of immune checkpoint blockade.” Proceedings of the National Academy of Sciences 117.44 (2020): 27509–27515.

29. Jin, Yueping, et al. “The diversity of gut microbiome is associated with favorable responses to anti–programmed death 1 immunotherapy in Chinese patients with NSCLC.” Journal of Thoracic Oncology 14.8 (2019): 1378–1389.

30. Tanoue, Takeshi, et al. “A defined commensal consortium elicits CD8+ T cells and anti-cancer immunity.” Nature 565.7741 (2019): 600–605.

31. Huang, Jumin, et al. “Ginseng polysaccharides alter the gut microbiota and kynurenine/tryptophan ratio, potentiating the antitumor effect of antiprogrammed cell death 1/programmed cell death ligand 1 (anti-PD-1/PD-L1) immunotherapy.” Gut (2021).

32. Lee SH, Cho SY, Yoon Y, Park C, Sohn J, Jeong JJ, Park H (2021) Bifidobacterium bifidum strains synergize with immune checkpoint inhibitors to reduce tumor burden in mice. Nat Microbiol 6(3):277–288.

33. Daillère, Romain, et al. “Enterococcus hirae and Barnesiella intestinihominis facilitate cyclophosphamide-induced therapeutic immunomodulatory effects.” Immunity 45.4 (2016): 931–943.

34. Vétizou, Marie, et al. “Anticancer immunotherapy by CTLA-4 blockade relies on the gut microbiota.” Science 350.6264 (2015): 1079–1084.

35. Pushalkar, Smruti, et al. “The pancreatic cancer microbiome promotes oncogenesis by induction of innate and adaptive immune suppression.” Cancer discovery 8.4 (2018): 403–416.

36. Griffin, Matthew E., et al. “Enterococcus peptidoglycan remodeling promotes checkpoint inhibitor cancer immunotherapy.” Science 373.6558 (2021): 1040–1046.

37. Wolf, Ashley R., et al. “Bioremediation of a common product of food processing by a human gut bacterium.” Cell host & microbe 26.4 (2019): 463–477.

38. Bui, Thi Phuong Nam, et al. “Production of butyrate from lysine and the Amadori product fructoselysine by a human gut commensal.” Nature communications 6.1 (2015): 1–10.

39. Yeoh, Yun Kit, et al. “Gut microbiota composition reflects disease severity and dysfunctional immune responses in patients with COVID-19.” Gut 70.4 (2021): 698–706.

40. Goloshchapov, Oleg V., et al. “Long-term impact of fecal transplantation in healthy volunteers.” BMC microbiology 19.1 (2019): 1–13.

41. Ott, Stephan J., et al. “Efficacy of sterile fecal filtrate transfer for treating patients with Clostridium difficile infection.” Gastroenterology 152.4 (2017): 799–811.

42. Sarrabayrouse, Guillaume, et al. “Mucosal microbial load in Crohn’s disease: A potential predictor of response to fecal microbiota transplantation.” EBioMedicine 51 (2020): 102611.

43. Sherry, Stephen, and Chunlin Xiao. “Ncbi sra toolkit technology for next generation sequence data.” Plant and Animal Genome XX Conference (January 14-18, 2012). Plant and Animal Genome. 2012.

44. Bolger, Anthony M., Marc Lohse, and Bjoern Usadel. “Trimmomatic: a flexible trimmer for Illumina sequence data.” Bioinformatics 30.15 (2014): 2114–2120.

45. Bushnell, Brian. BBMap: a fast, accurate, splice-aware aligner. No. LBNL-7065E. Lawrence Berkeley National Lab.(LBNL), Berkeley, CA (United States), 2014.

46. Olekhnovich, Evgenii I., et al. “Separation of donor and recipient microbial diversity allows determination of taxonomic and functional features of gut microbiota restructuring following fecal transplantation.” mSystems 6.4 (2021): e00811–21.

47. Beghini, Francesco, et al. “Integrating taxonomic, functional, and strain-level profiling of diverse microbial communities with bioBakery 3.” Elife 10 (2021): e65088.

48. Franzosa, Eric A., et al. “Species-level functional profiling of metagenomes and metatranscriptomes.” Nature methods 15.11 (2018): 962–968.

49. Kanehisa, Minoru, et al. “KEGG: new perspectives on genomes, pathways, diseases and drugs.” Nucleic acids research 45.D1 (2017): D353–D361.

50. Morton, James T., et al. “Establishing microbial composition measurement standards with reference frames.” Nature communications 10.1 (2019): 1–11.

51. Fedarko, Marcus W., et al. “Visualizing’ omic feature rankings and log-ratios using Qurro.” NAR genomics and bioinformatics 2.2 (2020): lqaa023.

52. Bolyen, Evan, et al. “Reproducible, interactive, scalable and extensible microbiome data science using QIIME 2.” Nature biotechnology 37.8 (2019): 852–857.

53. Olekhnovich, Evgenii I., et al. “MetaCherchant: analyzing genomic context of antibiotic resistance genes in gut microbiota.” Bioinformatics 34.3 (2018): 434–444.

54. Wood, Derrick E., Jennifer Lu, and Ben Langmead. “Improved metagenomic analysis with Kraken 2.” Genome biology 20.1 (2019): 1–13.

55. Oksanen, Jari, et al. “Package ‘ vegan’.” Community ecology package, version 2.9 (2013): 1–295.

56. Aitchison J. On criteria for measures of compositional difference. Mathematical Geol. 1992; 24(4):365–79. https://doi.org/10.1007/BF00891269.

57. Aitchison J. The one-hour course in compositional data analysis or compositional data analysis is simple In: Pawlowsky-Glahn V, editor. Proceedings of the III Annual Conference of the International Association for Mathematical Geology (vol.I). Barcelona, Spain: CIMNE: 1997. p. 3–35. ISBN 84-87867-97-9.

