## Supplementary figures and images for "Reproducible stool metagenomic biomarkers linked to the melanoma immunotherapy positive outcome"

### Supplelemntary Figure S1

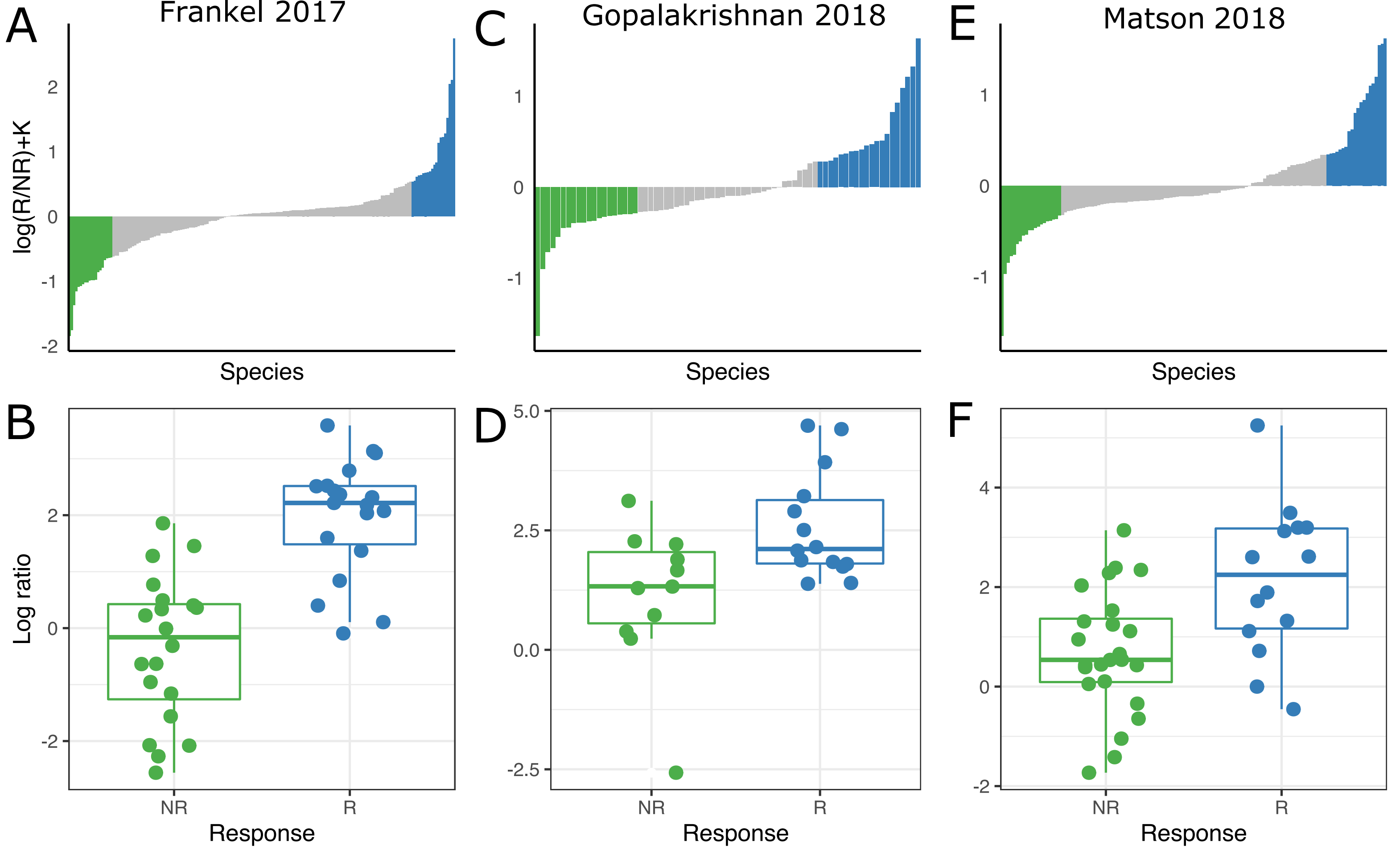

### Supplelemntary Figure S2

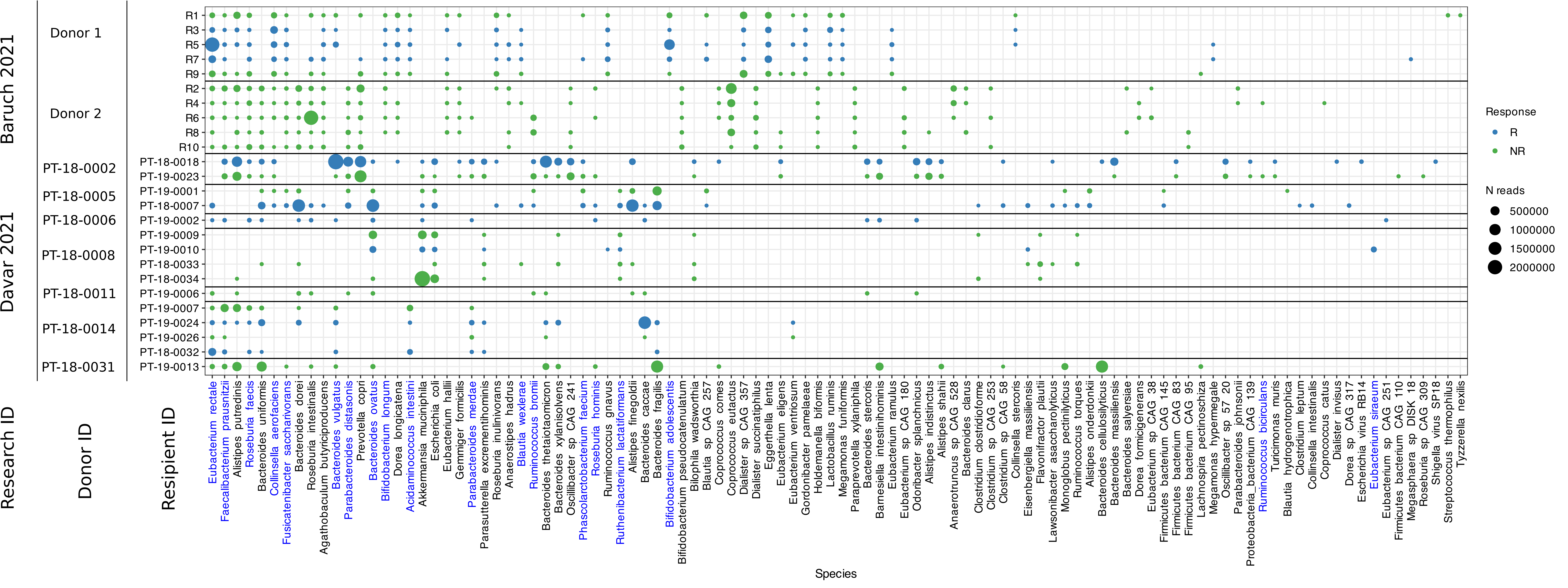

### Supplelemntary Figure S3

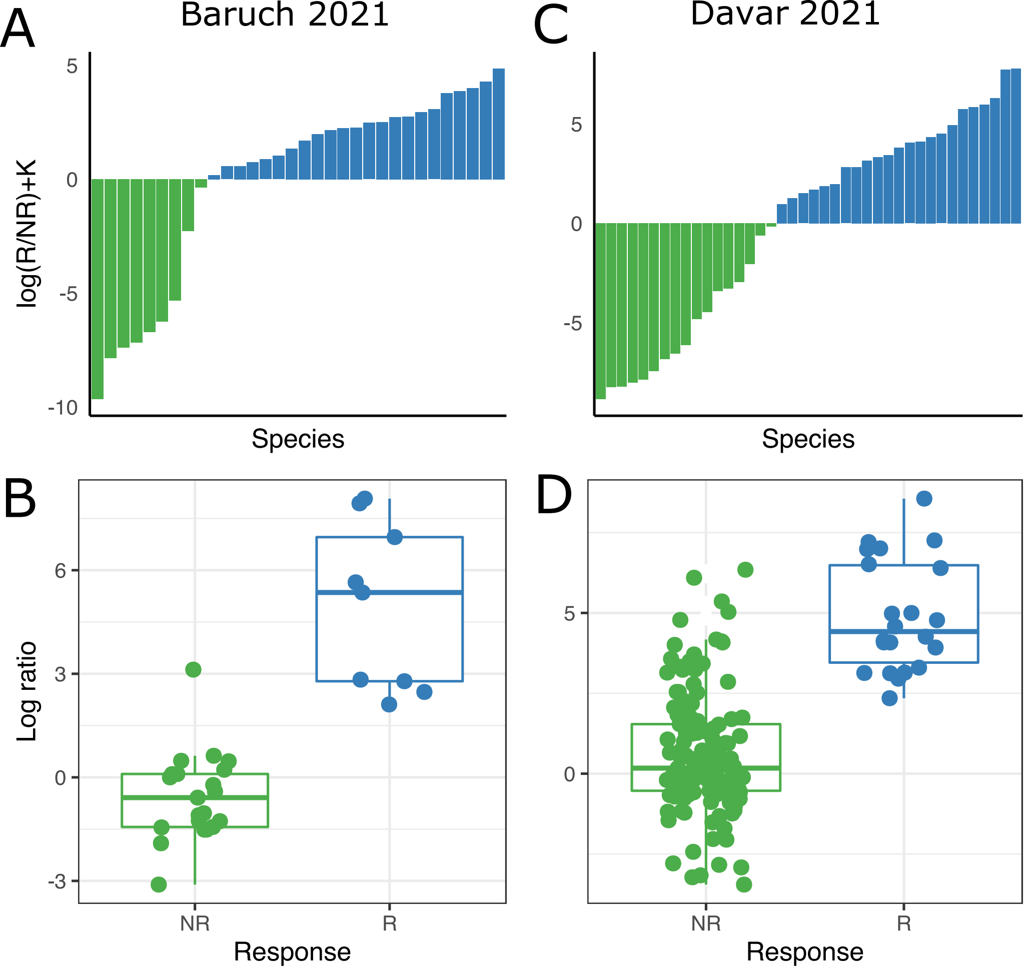
